# Low mammalian species richness is associated with Kyasanur Forest Disease outbreak risk in deforested landscapes in the Western Ghats, India

**DOI:** 10.1101/2021.04.08.439111

**Authors:** Michael G. Walsh, Rashmi Bhat, Venkatesh Nagarajan-Radha, Prakash Narayanan, Navya Vyas, Shailendra Sawleshwarkar, Chiranjay Mukhopadhyay

## Abstract

Kyasanur forest disease virus (KFDV) is a rapidly expanding tick-borne zoonotic virus with natural foci in the forested region of the Western Ghats of South India. The Western Ghats is one of the world’s most important biodiversity hotspots and, like many such areas of high biodiversity, is under significant pressure from anthropogenic landscape change. The current study sought to quantify mammalian species richness using ensemble models of the distributions of a sample of species extant in the Western Ghats and to explore its association with KFDV outbreaks, as well as the modifying effects of deforestation on this association. Species richness was quantified as a composite of individual species’ distributions, as derived from ensembles of boosted regression tree, random forest, and generalised additive models. Species richness was further adjusted for the potential biotic constraints of sympatric species. Both species richness and forest loss demonstrated strong positive associations with KFDV outbreaks, however forest loss substantially modified the association between species richness and outbreaks. High species richness was associated with increased KFDV risk but only in areas of low forest loss. In contrast, lower species richness was associated with increased KFDV risk in areas of greater forest loss. This relationship persisted when species richness was adjusted for biotic constraints at the taluk-level. In addition, the taluk-level species abundances of three monkey species (*Macaca radiata, Semnopithecus hypoleucus*, and *Semnopithecus priam*) were also associated with outbreaks. These results suggest that increased monitoring of wildlife in areas of significant habitat fragmentation may add considerably to critical knowledge gaps in KFDV epidemiology and infection ecology and should be incorporated into novel One Health surveillance development for the region. In addition, the inclusion of some primate species as sentinels of KFDV circulation into general wildlife surveillance architecture may add further value.

**Highlights:**

- Local mammalian species richness is estimated across the entire Western Ghats region
- Low species richness is associated with high KFDV risk in deforested landscapes
- This work identifies key landscapes for wildlife disease surveillance development

## Introduction

The sharing of space between wildlife and humans can be disruptive to both animals and people particularly in the context of altered habitat. Human pressure on wildlife influences species’ community composition, population movement and density, resource provisioning[1,2], and eco-immunology[3], which can subsequently alter pathogen circulation among wildlife hosts as well as introduce conduits that allow the movement of pathogens from reservoir hosts to novel human hosts[4], with potentially catastrophic effects[5–7]. As such, the growing wildlife-human interface requires both improved surveillance of pathogen circulation among wildlife and a greater understanding of the impact of anthropogenic landscape change on wildlife communities, their pathogens, and mechanisms for potential spillover of novel pathogens to humans. Importantly, as new wildlife-human interfaces emerge in close proximity to major transportation hubs, the potential for high-impact spillovers with the capacity for rapid regional or global dissemination is increasing[8].

Currently, there is a lack of zoonosis surveillance in wildlife globally, but the absence is particularly noteworthy in the world’s biodiversity hotspots that are undergoing rapid anthropogenic change. The Western Ghats region of South India is one of the world’s most important biodiversity hotspots[9] and also experiences significant pressure in the form of landscape change due to deforestation, commercial agricultural exploitation, and urbanisation[10]. The monsoon rainforests of this region support approximately 500 bird species, 225 reptile species, 219 amphibian species, and 133 mammal species[11,12]. Nevertheless, only 10% of the region’s 160,000 km^2^ area is formally protected[11]. As anthropogenic pressure increases in the region, conflict between many of these species, especially mammals, and humans has grown [13,14], facilitating exposure to novel pathogens. The general lack of wildlife pathogen surveillance across this rapidly changing region is compounded by an even greater absence of work examining the influence of community ecology on the processes of pathogen circulation and spillover events. The emerging tick-borne zoonotic virus, Kyasanur Forest disease virus (KFDV), simultaneously represents 1) a critical public health priority in the Western Ghats states because of its recent rapid expansion and the high mortality (2% - 10%) associated with its neurological and haematological complications [15–18], and 2) a model system for understanding viral spillover because of the extensive wildlife-human interface across the region[8]. Kyasanur Forest disease virus is a Flavivirus transmitted by several tick species, while the primary vector is the forest tick, *Haemaphysalis spinigera*. This species is found in high relative abundance in the region, has high viral prevalence, and feeds on many taxa of mammalian hosts including humans[19–21]. Humans typically are exposed to these ticks in the anthropogenic ecotones of forest fringe[22]. Outbreaks of KFDV have expanded from a single district in the state of Karnataka in the decade following the virus’ first identification in 1957 to an extensive region now comprising five states across South India[15,22]. Recent work has demonstrated a strong direct association between the expansion of KFDV outbreaks and the loss of native forest[23,24], which was further supported by phylogeographic analyses[25]. The study that reported the KFDV outbreak association with forest loss also noted that areas of high mammalian species richness presented greater risk[23]. While this previous study’s assessment of forest loss was of high quality, species richness was assessed more crudely using the IUCN-derived species’ range for each mammal species present across the region. As such, variation of species’ distributions within those ranges was not incorporated into the assessment of species richness, nor was there any assessment of potential interaction between sympatric species. Additionally, the associations between KFDV and individual species’ abundance and relative abundance were not previously explored. It is anticipated that further investigation of these aspects of mammalian biogeography and community ecology in KFDV outbreak hotspots will add considerable insight into the epidemiology of KFDV spillover while also providing an ecological evidence-base for developing municipal wildlife surveillance infrastructure at the taluk level (sub-district) across the Western Ghats states. The incorporation of wildlife monitoring and sampling into KFDV surveillance is particularly important since the investigation of infection in wildlife hosts has fallen off dramatically in recent decades, prior to which some mammalian susceptibility had been identified via serology but reservoir host competence was not established[15]. This decline in wildlife surveillance has resulted in a considerable knowledge gap in the fundamental infection ecology of KFDV, while also leaving any new efforts at building surveillance infrastructure uninformed with respect to what species should be sampled and where they should be sampled. Given the extensive forest loss in the region, delineating potential host communities across the spectrum of habitat fragmentation will be important in understanding how risk of spillover is modulated by wildlife communities. For example, as mentioned, the extent of viral competence is not known for most mammals of the Western Ghats and therefore identifying key reservoir hosts, or distinguishing between maintenance and amplification hosts, is not currently possible. As such, we remain largely ignorant of the importance of generalist species, which typically are more resilient to anthropogenic pressure, frequently dominate fragmented landscapes as biodiversity is lost, and often host zoonotic pathogens [26–28]. Conversely, the extent to which greater species richness in less fragmented landscapes may buffer against viral transmission (or against tick dispersal and feeding success) is also unknown. Therefore, there is an urgent need for targeted surveillance mechanisms to sample wildlife across heterogenous landscapes to fill these critical knowledge gaps and thereby ultimately develop a more sound approach to the control and prevention of this rapidly expanding tick-borne arbovirus.

The specific aims of the current study were as follows. First, we sought to estimate the species distributions of a sample of extant mammals of the Western Ghats (including endemic and non-endemic species) and use these to construct a more representative metric of species richness across the region. Second, we sought to adjust overall species richness, as well as individual species’ abundance and relative abundance for the biotic constraints of sympatry. Third, we sought to interrogate the association between species richness and KFDV outbreaks and the modification of this association by forest loss to develop much needed landscape epidemiology targets for improved surveillance of this expanding zoonotic arbovirus.

## Materials and Methods

### Data Sources

The Global Biodiversity Information Facility was used to obtain all direct observations of extant mammals in the Western Ghats[12] and Peninsular India between 1 January, 2010 and 1 November, 2020 (www.gbif.org). A total of 2826 mammals were observed over this period[29]. This comprised a total of 99 species observed across the region. However, habitat suitability was estimated only for those species with a sample size of at least 30 individuals, for a total of 24 species.

The KFDV outbreaks used in this investigation have been described in detail previously[23]. Briefly, two independent sources of outbreaks reported between 1 January, 2012 and 30 June, 2019 were used for model training and testing, respectively. The training data (47 outbreaks) were sourced from the ProMED-mail[30] electronic surveillance system and the testing data (39 outbreaks) comprised an independent set of laboratory-confirmed outbreaks described in the literature[23]. This allowed for external validation of KFDV outbreak models rather than partitioning data from a single source into training and testing.

Mean annual precipitation and mean annual temperature were obtained from the WorldClim Global Climate database at a resolution of 30 arc seconds (∼ 1 km) [31]. The rasters derived from the WorldClim database were mean measurements between the period 1950 to 2000, and thus represent estimates of climate rather than weather. Some regions represented by the WorldClim database have sparse weather stations contributing to climate estimates. However, India’s extensive network of weather stations make its local contributions to decadal climate interpolation more representative than many other large countries[32].

The Priestley-Taylor α coefficient (P-Tα) is the ratio of actual evapotranspiration to potential evapotranspiration and was included here as a metric for water-soil balance[33,34]. In contrast to solar energy input alone, the P-Tα represents water availability in the soil as a function of the local vegetation’s water requirements and is therefore a more robust estimate of soil-water balance. A 30 arc seconds resolution raster of P-Tα was obtained from the Consultative Group for International Agricultural Research (CGIAR) Consortium for Spatial Information. The ratio ranges from 0 (extreme water stress) to 1 (no water stress)[35].

Forest cover was first acquired from the Global Land Cover Climatology MODIS-based data product, which corrects temporally aggregated 10-year land cover types by weighting each land cover type by their annual uncertainty and then validates the weighted composite against the System for Terrestrial Ecosystem Parameterization[36]. The product used here for forest cover represents the period 2001-2010 at a resolution of 15 arc seconds (∼ 500 m).

Landsat data compiled at 1 arc second (∼ 30 m) resolution by the Global Forest Change project was used to quantify forest loss between 2000 and 2012[37]. Landsat imagery was processed using a stratified random sampling validation procedure, with 99.6% accuracy in global settings and 99.5% accuracy in tropical settings[37]. Individual tiles were merged across the region of the Western Ghats and a new quantile raster constructed to represent deciles of forest loss.

An established validated metric for the human footprint (HFP) was used to control for spatial background sampling as described below in the analysis section. The metric was acquired from the Socioeconomic Data and Applications Center (SEDAC) repository[38] at a resolution of 30 arc seconds and has been described in detail previously[39]. Briefly, HFP comprises 2 levels of classification. First, an index of human influence was determined based on eight domains: 1) population density, distance to 2) roads, 3) railway lines, 4) navigable rivers, and 5) coastlines, 6) degree of night-time artificial light, 7) urban vs rural location, and 8) land cover. These domains were then scored to derive the human influence index (HII), ranging from 0 (no human impact) to 64 (greatest human impact). The ratio of the range of minimum and maximum HII in the local terrestrial biome to the range of minimum and maximum HII across all biomes represents the final HFP metric and is expressed as a percentage[39].

Because KFDV outbreaks disproportionally affect marginalised communities with limited access to health care, this investigation adjusted for potential reporting bias using a measure of the distribution of health system performance as an indication of the local capacity to detect outbreaks (see modelling description below). The infant mortality ratio (IMR) was chosen as a proxy for health system performance because this has been verified as a robust indicator of health infrastructure and health system performance and used extensively to compare health service delivery[40,41]. The IMR strongly correlates with disability-adjusted life expectancy (DALE), the Human Development Index (HDI), and the Inequality-Adjusted Human Development Index (IHDI). As such, the IMR is an important indicator of structural issues that affect health care access and delivery, such as economic development, general living conditions, social well-being, and the quality of the environment[40,42]. The raster of the IMR was obtained from SEDAC[43].

### Data Analysis

#### Species distribution modelling

The landscape suitability of each of the 24 mammal species was estimated using an ensemble approach comprising two machine learning methods (boosted regression trees (BRT) and random forests (RF)) and generalised additive models (GAM)). The machine learning frameworks, BRT and RF, are powerful approaches to improving standard decision trees by iteratively growing many trees and amalgamating the results. As the algorithms partition the data space according to rules that optimise homogeneity among predictors and a response (e.g. species presence), many decision trees are grown and combined, resulting in optimised decision trees that reduce overfitting and can capture complex interactions between the predictors [44–47]. There are two key differences in these machine learning approaches. First, for each decision tree partition in RF, only a random subset of predictors is selected from the set of all predictors, which decorrelates the trees and reduces overfitting. Second, rather than decorrelating trees based on the sampling of subsets as with RF, BRT instead reduces overfitting by growing trees sequentially and learning from previously grown trees. In contrast, the GAM framework allows for nonlinear relationships between outcomes and covariates by fitting multiple basis functions for smoothed covariates [48,49]. Five-fold cross-validation was used to fit each model under the three distinct modelling frameworks (BRT, RF, and GAM). Mean annual temperature, P-T α, and forest cover were included as landscape features at 30 arc seconds resolution in all models. For each species, each of the three models describing their landscape suitability was evaluated according to performance, based on the area under the receiver operating characteristic curve (AUC), and fit, based on the model deviance. Subsequently, an ensemble landscape suitability was computed from the three model classes using their weighted mean, with weights based on AUC [50]. To adjust for potential spatial sampling bias in the GBIF database, background points were sampled proportional to the human footprint as a proxy for landscape accessibility. The estimates of each species’ distribution as derived from these ensembles were then summed across all species to estimate local species richness across the Western Ghats region. This estimate as well as an additional estimate adjusted for the biotic constraints of sympatric species (see description below) were subsequently used as more appropriate metrics for mammalian species richness and the evaluation of its influence on KFDV outbreaks. As an additional sensitivity analysis to test whether associations with species richness may be influenced by the limited regional sample of mammalian species used to estimate species richness, an alternate India-wide sample (number of observations = 5833, number of species = 30) was used to re-estimate individual species habitat suitability and then species richness. The sdm package[50] in the R platform[51] was used for fitting each model and the derivation of the three-model ensembles to each species.

#### Point process modelling

The KFDV outbreaks were fitted as an inhomogeneous Poisson point process[52,53]. Under this model framework the spatial dependence of the outbreaks’ distribution can be determined and evaluated with respect to specific landscape features. The background points used in these models were sampled proportional to IMR, as described above, to control for potential outbreak reporting bias. Previous work demonstrated a strong association between KFDV outbreaks and both forest loss and mammalian species richness[23]. However, the association with the latter was 1) based on a crude representation of species richness that did not account for local heterogeneity of individual species’ distributions and 2) could not be used to evaluate the potential effect modification of species richness by forest loss. The current investigation provides this critical missing component by incorporating a more suitable metric of species richness based on the individual estimated species’ distributions described above and subsequently interrogating the interaction between species richness and forest loss. In addition, to account for the important climate features that were incorporated in the previous work, precipitation and temperature were also included in the current work. Finally, the possibility of confounding of the associations between KFDV outbreaks and species richness and forest loss by alternative forms of human influence in the landscape was assessed by including HFP as a covariate in an additional model. Therefore, the suite of multiple inhomogeneous Poisson models comprised the newly-quantified mammalian species richness metric described above, forest loss, an interaction term for species richness and forest loss, mean annual precipitation, mean annual temperature, and HFP all aggregated up to 2.5 arc minutes. The landscape features included in the multiple inhomogeneous Poisson models were not highly correlated so multicollinearity in the models was not of concern (all values of the Pearson’s r were < 0.5). Model fit was assessed using the Akaike information criterion (AIC) and model performance was tested against an independent, laboratory-confirmed set of outbreak data, as described above. Performance was evaluated using the AUC. The spatstat R package was used to fit the point process models[54].

#### Taluk-level modelling

Species richness was further adjusted for, and individual species abundance and relative abundance were computed based on, the potential modulating effects of sympatric species at the scale of the taluk. Taluks are 3rd-level, subdistrict administration units in India and are small enough to reasonably represent shared space between sympatric species, while simultaneously large enough to delineate the minimal municipal infrastructure (i.e. delineating the most local focus) used in the Western Ghats states for the implementation of animal and human disease surveillance. Specifically, biotic constraints were applied to the estimates of each species distribution according to the spatially-explicit species assemblage modelling (SESAM) framework[55–57]. Under this framework, first the landscape suitability for each species was estimated according to the ensemble method described above. Second, species richness was calculated as the sum of the individual species distributions, again using the method described above. Third, the biotic constraint was applied whereby each species is evaluated with respect to all other species present within each taluk via the probability ranking rule[56,58] to determine whether a given species should be retained within, or excluded from, each taluk “community”. The third step ranks each species from highest to lowest predicted habitat suitability based on each species’ suitability obtained in the first step. Then, species are selected for inclusion beginning with the species with highest suitability and proceeding down the catalogue of ranked species until the sum of selected species is equal to the expected species richness value for each location calculated in the second step. This provides an estimate of each species’ presence (or absence) at the taluk level given the potential biotic constraints of the other species within the taluk. For those species thusly identified as present within the taluk community, all 1 km^2^ pixels exceeding the true skill statistic (TSS)[59] of their ensemble landscape suitability estimate were designated present (1 = present, 0 = otherwise) and summed across all pixels within the taluk. This yielded a taluk-level estimate of species abundance for each species under consideration. Similarly, the taluk-level relative abundance for each species was quantified by dividing each individual species abundance by the total abundance of all species identified as present within each taluk under the SESAM framework. Finally, a new estimate of species richness was computed by summing all the species identified as present within each taluk under the SESAM framework. This systematic approach thus yielded taluk-level community estimates of each species’ abundance and relative abundance, as well as an updated estimate of species richness, all adjusted for the biotic constraints of sympatric species. The SESAM framework, including the probability ranking rule was implemented using the ecospat package in R[57].

Integrated nested Laplace approximation (INLA) models[60] were used to estimate the association between KFDV outbreaks and the biotically constrained estimate of mammalian species richness and its interaction with forest loss at the level of the taluk, as well as associations between KFDV outbreaks and individual species abundance and relative abundance to identify potential species that may be good candidates for initial wildlife surveillance. While the latter may represent species that are important as reservoir hosts, we did not attempt to assess species’ roles as hosts in the current study since we did not measure their competence for KFDV. Instead, individual species associations were investigated simply to identify whether certain species may be useful sentinels of KFDV outbreaks. The binomial likelihood family was used for the INLA models with spatial autocorrelation estimated using random effects with Besag–York–Mollie priors. Under the binomial family, taluks were modelled as either having experienced a KFDV outbreak under the period of study (outbreak positive) or not (outbreak negative). This approach was taken since the majority of taluks in the region did not experience outbreaks, and, among those that did, most only experienced one or two outbreaks. Nevertheless, to evaluate the effect of different distribution families, we also fitted these models with the zero-inflated Poisson family as a sensitivity analysis. Since the KFDV outbreaks were necessarily aggregated by taluk, to increase the outbreak sample size available for taluk-level modelling, the independent datasets described above for the point process models (PPMs) were combined for the INLA models. Since the primary objective was estimation rather than prediction so as to infer associations between KFDV outbreaks and species richness, this was deemed an acceptable trade-off. The fit of INLA models was evaluated using the Watanabe-Akaike information criterion (WAIC). The INLA models were fit using the inla package in R (www.r-inla.org) [60].

## Results

The distribution of mammalian species richness, estimated as the sum of the sample of individual species’ distributions, is presented in Figure 1 juxtaposed with the distribution of KFDV outbreaks. The individual species distribution models generally performed well (Supplementary material, S1 Table 1). The distribution of each species’ landscape suitability is presented in S2 Figure 1 and S3 Figure 2. Both forest loss and the new quantification of mammalian species richness based on individual species’ landscape suitability alone were very strongly associated with KFDV outbreaks at local scale (Table 1). Strong associations were apparent whether these features were considered alone in the crude bivariate PPMs (Table 1A), or together in the multiple PPMs (Table 1B, 1C, and 1D). Critically, while species richness demonstrated a similar association with KFDV outbreaks to that shown previously[23], the current investigation identified significant and substantial interaction between species richness and forest loss. As such, increasing mammalian species richness was crudely associated with increasing KFDV outbreak occurrence. However, when species richness was considered in concert with forest loss, the KFDV association with the former was modified by the latter (coefficient = -0.41). This relationship was maintained in the final model (Table 1C), which incorporated the climate features and was a markedly better fit (AIC = -36.34 vs. -22.80) although performance was slightly diminished (AUC = 0.74 vs. 0.78). Finally, confounding of these associations by HFP was not supported since the regression coefficients were unaltered by the inclusion of HFP, HFP itself was not associated with outbreaks, and the model overall was a poorer fit to the data (Table 1D). The distribution of KFDV outbreak risk across the Western Ghats based on the best fitting model (Table 1C) is presented in Figure 2. The individual species distribution models and projected suitability using the alternate country-wide sample are presented in S4 Table 2 and S5 Figure3 and S6 Figure 4. There was no marked difference in the strength or direction of the association between KFDV outbreak risk and species richness or its interaction with forest loss in the sensitivity analysis using the India-wide sample, however the model based on the latter demonstrated both poorer fit and performance (S7 Table 3, S8 Figure 5), so the original model based on the regional sample was retained as the primary model for this analysis.

**Figure 1.**
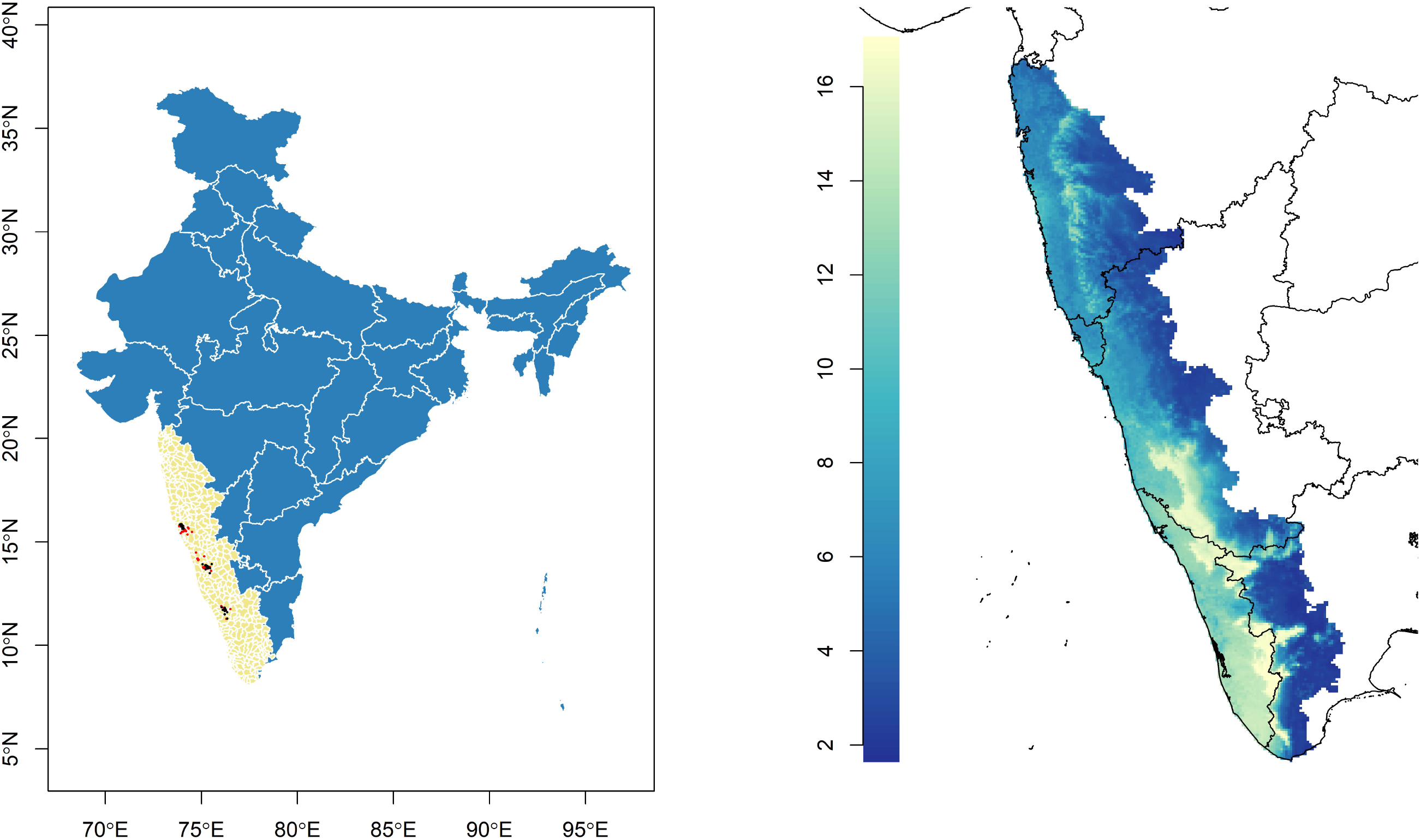
The distribution of taluks (beige) across the general region of the Western Ghats comprises five states (Maharashtra, Goa, Karnataka, Kerala, and Tamil Nadu) in Peninsular India (left panel). Kyasanur Forest Disease virus outbreaks captured via ProMED-mail (red) and independently sourced (black) are superimposed over taluks. The distribution of mammalian species richness as estimated from a sample of Western Ghats species from 2010 to 2020 is presented in the right panel.

**Table 1.** Unadjusted (A) and adjusted (B, C, and D) regression coefficients and 95% confidence intervals for the associations between Kyasanur Forest disease virus outbreaks and landscape features as derived from inhomogeneous Poisson point process models (PPMs) of the outbreaks. Coefficients in A represent crude associations from bivariate PPMs, whereas those in B, C, and D represent adjusted associations wherein each landscape feature is adjusted for all other features included in the multiple PPMs. The Akaike information criterion (AIC) and area under the receiver operating characteristic curve (AUC) show model fit and performance, respectively.

| Landscape features | Coefficient | 95% confidence interval | AIC | AUC |
| --- | --- | --- | --- | --- |
| **A. Bivariate PPM** |  |  |  |  |
| Mammal richness | 0.08 | 0.02 – 0.15 | 31.79 | 0.62 |
| Forest loss (deciles) | 1.22 | 0.85 – 1.59 | 27.14 | 0.76 |
| Annual precipitation (100 mm) | 0.06 | 0.04 - 0.08 | 28.64 | 0.63 |
| Annual temperature (Celsius) | -0.02 | -0.03 – -0.01 | 56.88 | 0.53 |
| **B. Multiple PPM – without climate** |  |  |  |  |
| Mammal richness | 0.59 | 0.37 – 0.82 | -22.80 | 0.78 |
| Forest loss (deciles) | 5.21 | 3.94 – 6.48 |  |  |
| Mammal richness:forest loss interaction | -0.41 | -0.54 – -0.28 |  |  |
| **C. Multiple PPM – with climate** |  |  |  |  |
| Mammal richness | 0.39 | 0.12 – 0.65 | -36.34 | 0.74 |
| Forest loss (deciles) | 4.82 | 3.46 – 6.19 |  |  |
| Mammal richness:forest loss interaction | -0.37 | -0.51 – -0.23 |  |  |
| Annual precipitation (100 mm) | 0.05 | 0.02 – 0.08 |  |  |
| Annual temperature (Celsius) | -0.03 | -0.05 – -0.01 |  |  |
| **D. Multiple PPM – with the human footprint** |  |  |  |  |
| Mammal richness | 0.39 | 0.12 – 0.66 | -35.34 | 0.75 |
| Forest loss (deciles) | 4.86 | 3.47 – 6.24 |  |  |
| Mammal richness:forest loss interaction | -0.37 | -0.51 – -0.23 |  |  |
| Annual precipitation (100 mm) | 0.05 | 0.02 – 0.08 |  |  |
| Annual temperature (Celsius) | -0.03 | -0.05 – -0.01 |  |  |
| Human footprint (%) | -0.02 | -0.05 – 0.02 |  |  |

**Figure 2.**
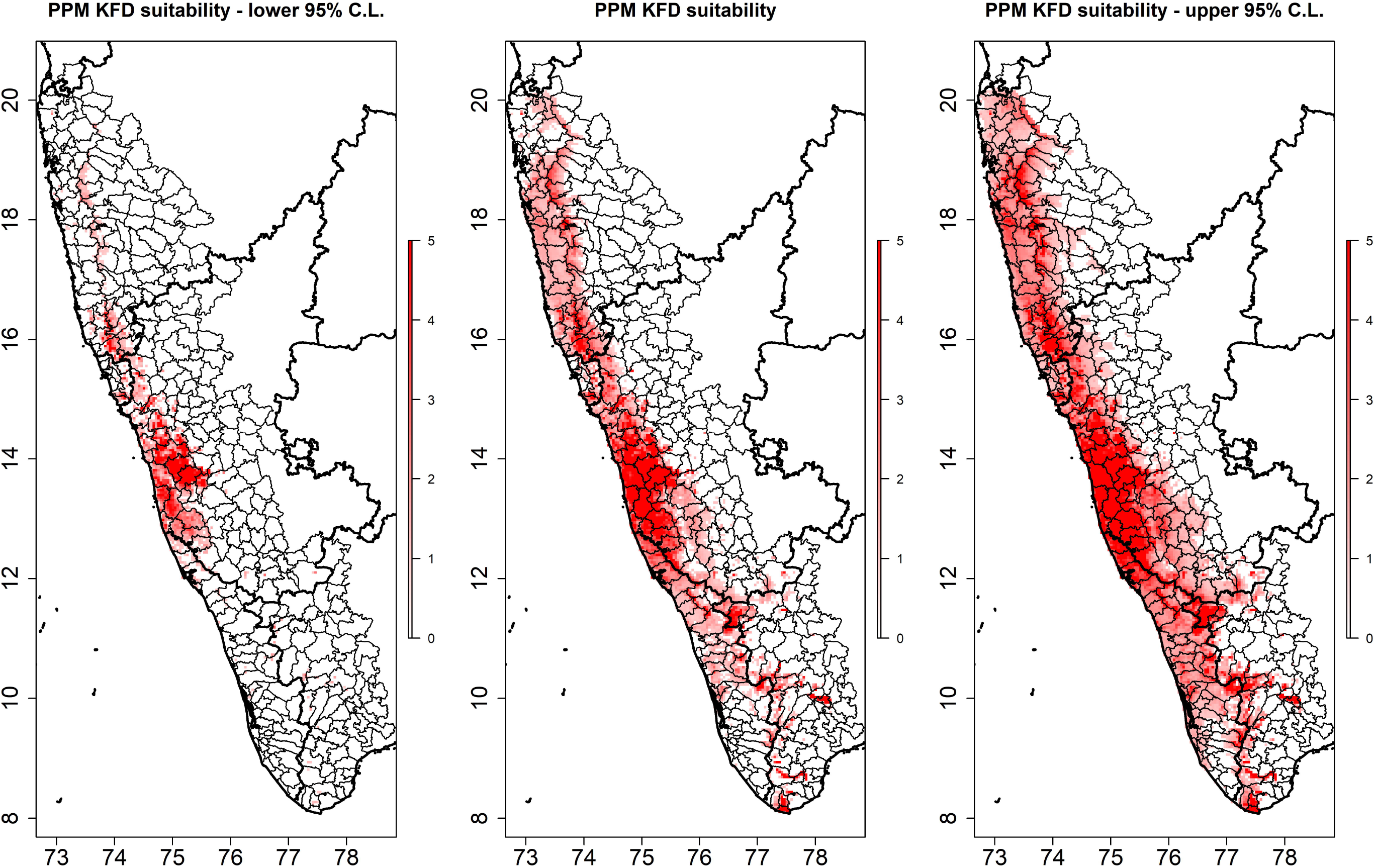
The distribution of Kyasanur Forest Disease virus outbreak risk derived from the inhomogeneous Poisson model is presented in the centre panel with the lower and upper 95% confidence limits to the left and right, respectively.

The newly quantified metric for taluk-level species richness based on the SESAM framework is presented in Figure 3. In comparison to the best-fitting high-resolution PPM model (Table 1C), the taluk-level INLA model showed the same strong interaction between forest loss and mammal species richness, even as the latter was adjusted for biotic constraint under the SESAM framework and scaled up to taluk (Table 2, Figure 4). Further sensitivity analysis showed that these associations all persisted when the distribution family was specified as zero-inflated Poisson (S9 Table 4 and S10 Figure 6), although temperature was no longer associated with outbreak occurrence. Finally, as with the PPMs described above, the associations persisted (S11 Table 5) when species richness was estimated based on the alternate India-wide sample (S12 Figure 7) although the model overall was a poorer fit. Based on taluk-level outbreak risk (Figure 4), we identified the 50^th^ percentile of risk where deforestation and species richness optimally converge for targeted selection for a state-administered (e.g. Karnataka) animal surveillance program (Figure 5). In addition to the broad metric of species richness, we also explored the associations between KFDV outbreaks and each individual species’ abundance and relative abundance to determine if any particular species may prove useful as a sentinel surveillance target. Despite several species demonstrating significant individual associations with KFDV occurrence at fine scale based on their estimated landscape suitability (S13 Table 6), most species did not demonstrate associations with outbreaks at the level of the taluk (S14 Table 7). However, there were a few notable exceptions. Species abundance of the primates *Macaca radiata, Semnopithecus hypoleucos* and *Semnopithecus priam*, were all associated with increased KFDV occurrence at the taluk level. Species relative abundance was also associated with KFDV outbreaks for *S. hypoleucos*, but not for any other primate species. The distributions of these species’ abundances and relative abundances per taluk are presented in S15 Figure 8. Finally, both the abundance and relative abundance of *Panthera pardus*, and the abundance of *Axis axis*, were also associated with increased KFDV outbreak occurrence.

**Figure 3.**
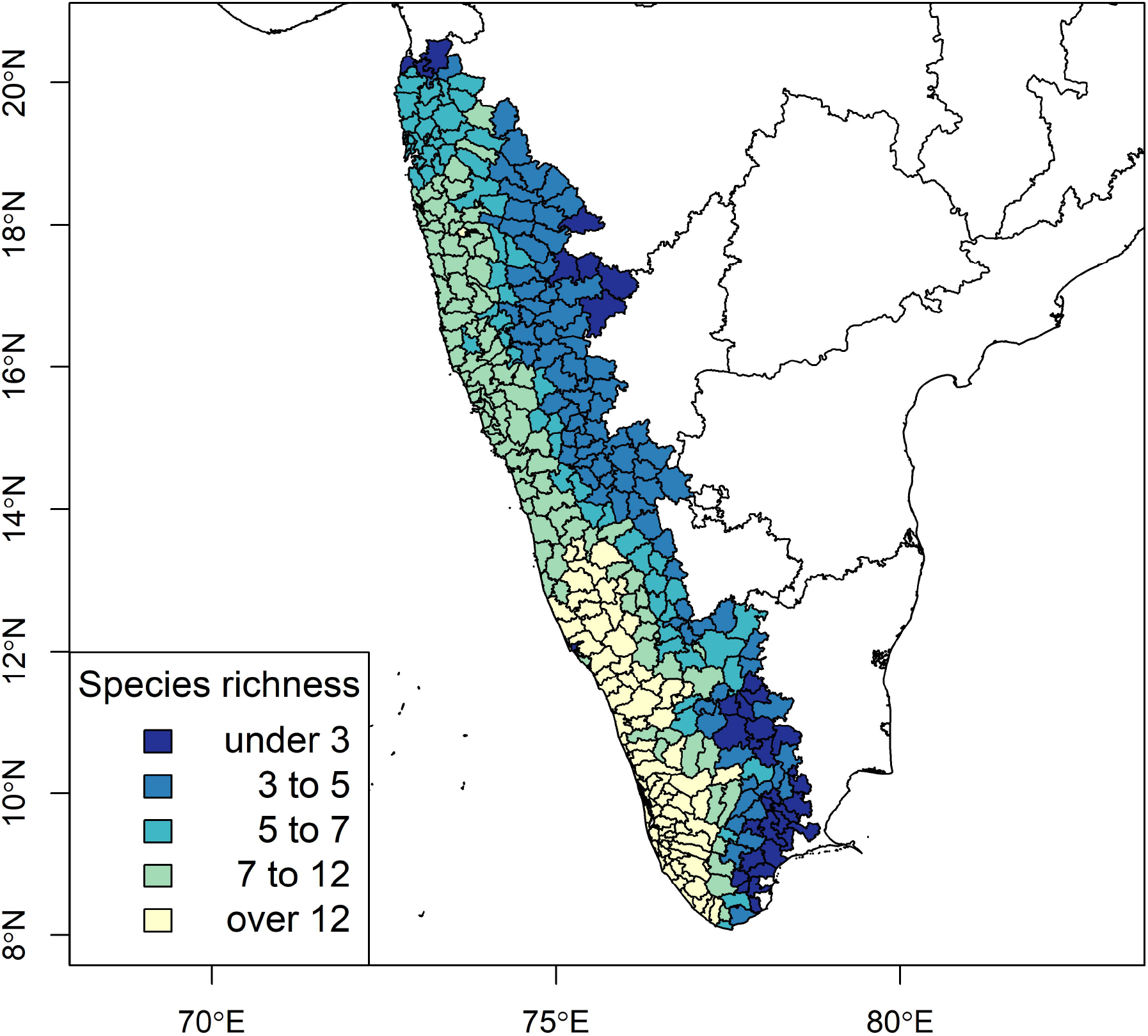
The adjusted distribution of mammalian species richness as estimated from a sample of Western Ghats species. Species richness was adjusted using the SESAM framework to introduce biotic constraints to adjust the presence of each species for the presence of other species within each taluk.

**Table 2.**
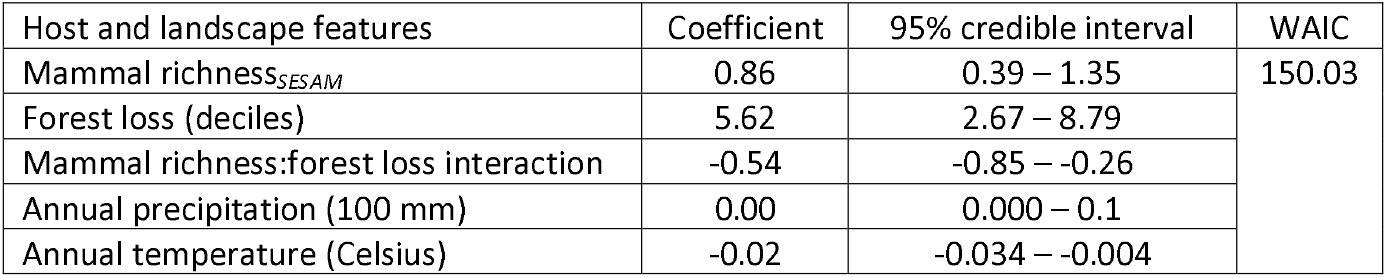
Taluk-level integrated nested Laplace approximation model (binomial family) of Kyasanur Forest Disease virus outbreaks. Mammal species richness was adjusted for the biotic constraints of sympatric species using the SESAM framework. The Watanabe-Akaike information criterion (WAIC) was used to assess model fit.

**Figure 4.**
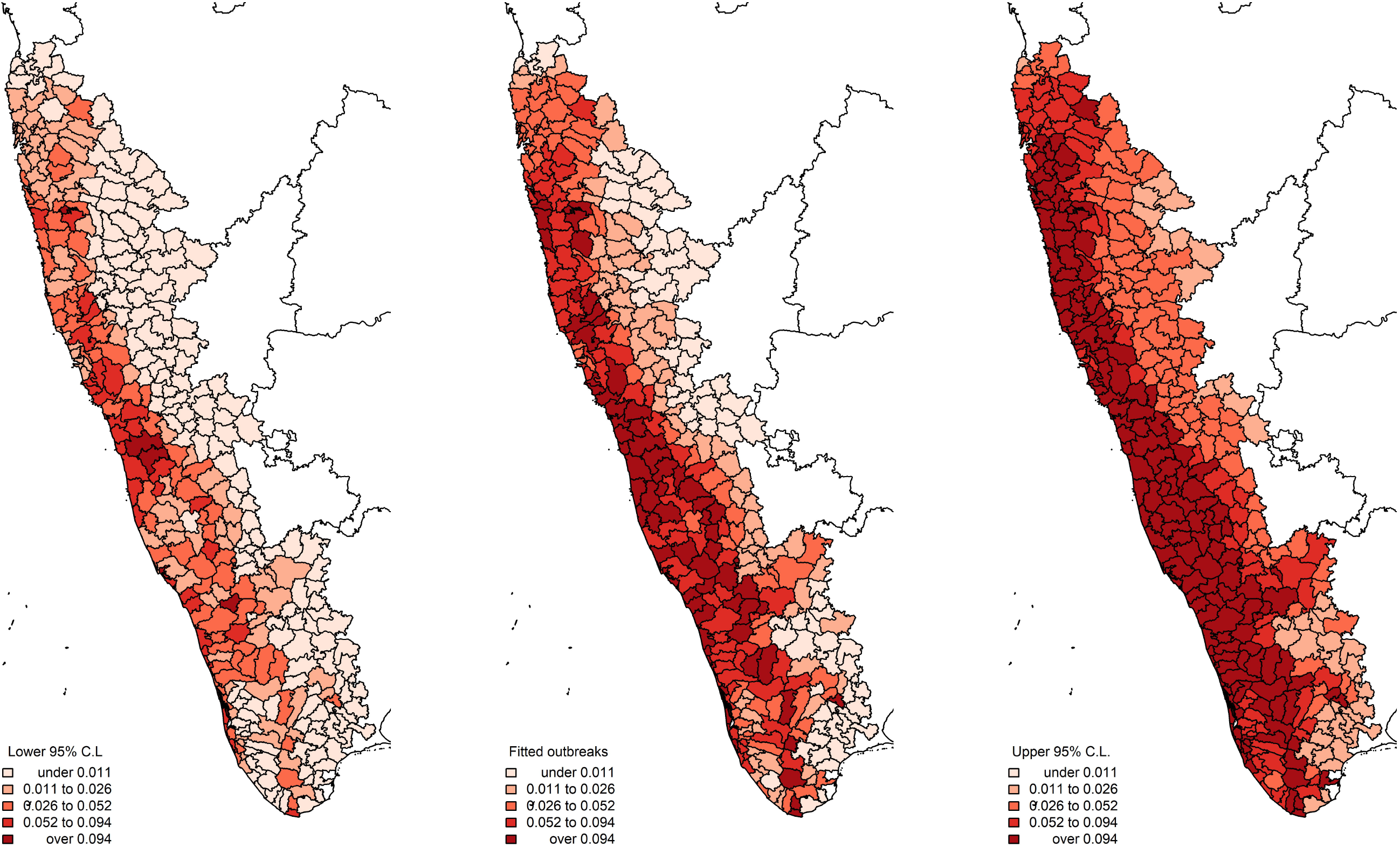
The distribution of Kyasanur Forest Disease virus outbreak risk derived from the integrated nested Laplace approximation model (binomial family) is presented in the centre panel with the lower and upper 95% credible limits to the left and right, respectively.

**Figure 5.**
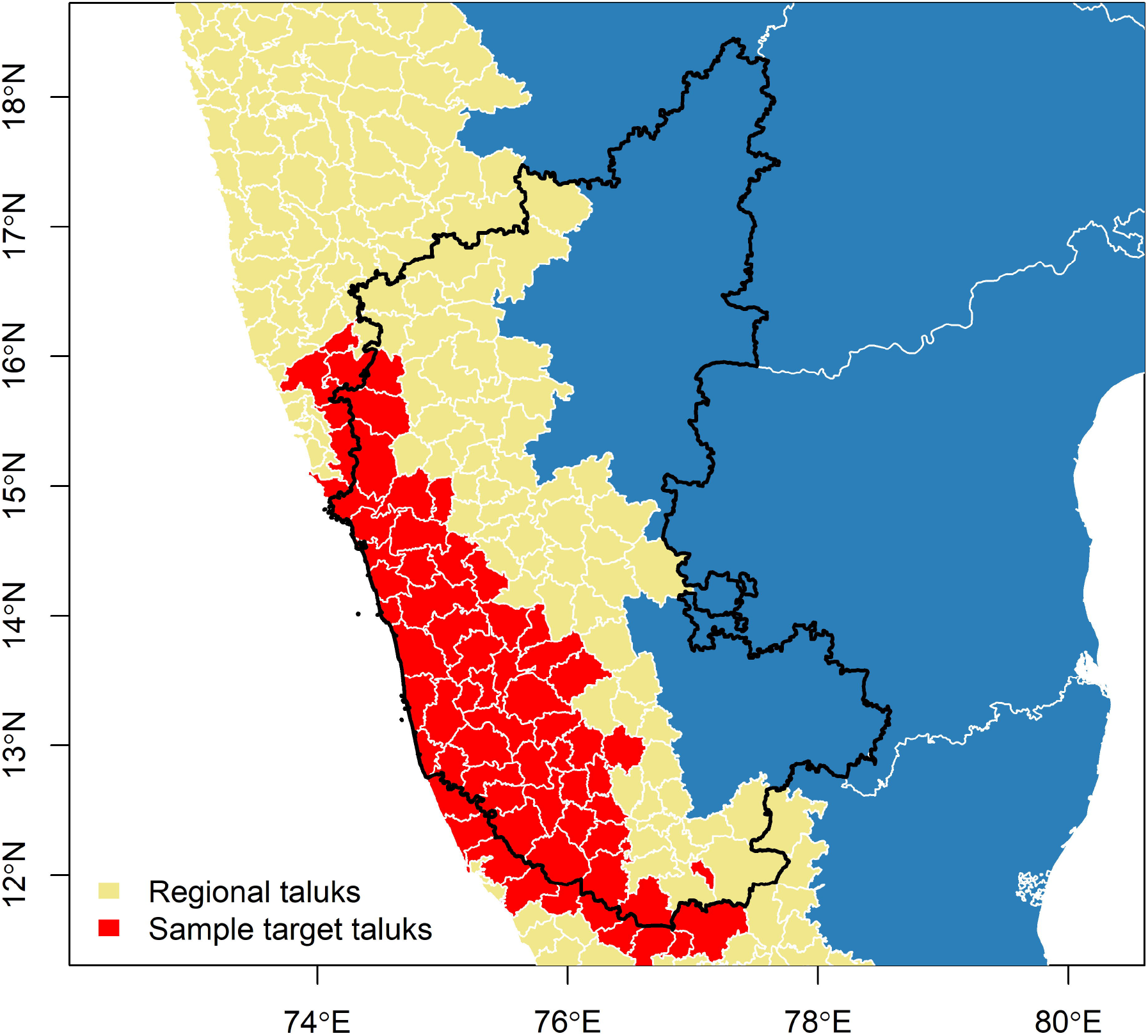
Taluks identified as priority targets (red) for implementing animal-human surveillance in Karnataka state. Having located priority taluks, a sample of these can be taken either by random draw or proportional to the distribution of key landscape features, such as species richness and/or forest loss. Once the taluks are selected, One Health surveillance infrastructure can be developed by 1) working in concert with the forest department to select sampling grids from within the sampled taluks for the monitoring and sampling of wildlife, 2) working in concert with animal husbandry and veterinary services to generate random samples of farms/livestock holders from animal censuses to monitor and sample cattle, and 3) working in concert with primary healthcare centres to improve human case detection and care delivery, conduct serosurveys, and implement prevention outreach.

## Discussion

This is the first study to attempt a general estimate of mammalian species richness across the Western Ghats region based on ensemble modelling of the landscape suitability of several extant species, and to extend this by adding biotic constraints of sympatric species to the estimates of species richness, as well as individual species abundance and relative abundance. This is also the first study to demonstrate negative effect modification of the association between mammalian species richness and KFDV outbreak occurrence by forest loss such that higher species richness was associated with increased KFDV outbreak risk where deforestation was minimal, but in areas of increasing forest loss lower species richness was associated with increased risk. This association was identified at both fine scale and at the level of the taluk. Finally, three monkey species were identified (*M. radiata, S. hypoleucus*, and *S. priam*) whose taluk-level species abundances were indicators of outbreaks. Taken together, these findings have identified important interaction between landscape degradation and mammalian species richness that may influence the ecology of this tick-borne virus in its natural foci. The fact that this relationship held at the taluk level even after controlling for biotic constraints indicates that enhanced monitoring of wildlife and human populations at the level of the taluk, which is well-suited to the municipal infrastructure of current human and animal health systems in the Western Ghats states, may offer promise as the focus of novel One Health surveillance development. In addition, the incorporation of some primate species as sentinels of KFDV circulation into general wildlife surveillance may add further value to such a One Health program.

Previous work identified increased risk of KFDV outbreaks associated with both forest loss and high mammalian species richness[23]. However, the latter association was difficult to interpret due to the fairly crude measure of species richness, which was simply a composite of IUCN mammal species’ ranges across the region. Moreover, this work was not able to assess interaction between forest loss and mammalian biodiversity. The positive association between KFDV outbreak risk and species richness shown in that study was reasonable given the expectation of greater exposure to potential KFDV hosts and their tick vectors following increasing incursion into areas of high species richness. Nevertheless, it is also of particular interest to identify how habitat loss may affect the community ecology of mammalian hosts in ways that could influence KFDV spillover to humans. Therefore, better metrics of species richness were required across the region to more appropriately evaluate the modifying effect of forest loss on the association between species richness and KFDV outbreaks. The results presented in this current investigation indicated that the relationship between species richness and forest loss is indeed more nuanced. As shown previously[23], increased species richness was associated with increased KFDV outbreak risk but in the current study this relationship only held in landscapes with minimal forest loss. In contrast, as forest loss increased, lower species richness was associated with increased risk. While this finding allows no definitive assertion regarding the nature of KFDV circulation among mammalian hosts, such as the relative importance of the dilution effect to transmission dynamics[61], it does provide important evidence that KFDV circulation among reservoir hosts persists, and spillover to humans continues, in human-altered landscapes exhibiting lower biodiversity, which is consistent with previous evidence showing clear pathways from habitat fragmentation and biodiversity loss to increased spillover[1,62]. Despite the current study’s inability to make specific claims about the role of any individual species in the viral maintenance and infection ecology of KFDV, the findings do suggest that, in general, species resilient to anthropogenic landscape change may be most relevant to the expansion of KFDV spillover across the region. This finding closely parallels recent work demonstrating similar relationships more globally for other zoonotic pathogens[26,27]. Moreover, the work may help to address a critical surveillance gap currently impeding a fuller understanding of KFDV infection ecology. Following the initial identification of KFDV in 1957, there was a rapid but limited flurry of research attempting to identify wildlife reservoirs for KFDV in the Bandipur Forest range[15]. Several mammalian species were identified as hosts, but these studies were not definitive with respect to KFDV infection ecology as many relied on serology, rather than measuring host competence, and many of the studies employed limited sampling strategies[15]. Subsequently, wildlife investigations all but stopped by the early 1980s before KFDV reservoirs could be established. Therefore, the current study could have important implications for reinstating widespread wildlife surveillance of KFDV in the Western Ghats with a focus on synanthropic species.

It is important to caution against interpretation of these findings to infer specific dynamics of infection ecology at either the level of the community or individual species because infection competence was not assessed in any of the mammals under investigation nor was interspecific interaction among species directly observed at species’ locations. However, the results do provide some of the most actionable evidence to date for targeted, landscape-designed wildlife surveillance across the region. Specifically, these findings can help to frame a landscape-based cluster sampling design for the monitoring of wildlife distribution and movement in response to human induced landscape change in concert with pathogen surveillance in animals and humans (Figure 5). Critically, the fact that this study’s findings were consistent at the level of the taluk indicates that the existing municipal infrastructure for animal and human health can be used as the necessary scaffolding for novel One Health surveillance architecture and service delivery. Moreover, the taluk-level abundance of bonnet macaques (*M. radiata*) and grey langurs (*Semnopithecus spp*.), which have previously been identified as highly susceptible to KFDV[63–65], were also associated with outbreaks. As mentioned above, the current study can make no claims regarding the roles of individual species in the infection ecology of KFDV, but the associations with these primate species’ abundance within taluks indicates that these species may provide useful sentinels under a One Health surveillance framework. In addition, leopard (*P. pardus*) abundance and relative abundance and chital (*A. axis*) abundance were also associated with outbreak risk. While neither of these species have been shown to be susceptible to KFDV, both are known to host KFDV-relevant ticks and distribute these vectors in the landscape[66,67]. Therefore, their associations with increased KFDV outbreak occurrence in the current study merits a closer look at the role these species could play as tick distributors, particularly since both of these species are highly adaptive to anthropogenic landscapes. As such leopards and chital could have repercussions for human exposure to KFDV vectors, their unlikely role as important KFDV hosts notwithstanding.

Further to the cautionary interpretation around KFDV infection ecology encouraged above, some further description of this study’s limitations is provided. First, as previously mentioned direct observation of interspecific interaction was not possible under the current investigation, which precludes the ability to make specific claims about the true nature and influence of interaction among communities at various scales across the region. The findings reported here will need to be verified by systematically conducted field investigations of directly observed interspecific interaction at multiple scales. Second, sufficient observations were not available for each extant mammal species of the Western Ghats and so the new quantification of species richness presented here, although much improved over previous estimates, is based on only a sample of Western Ghats mammals and therefore remains a proxy for true species richness. Third, spatial biases may affect the distribution of species observations, as well as the reporting of KFDV outbreaks, across the region. To control for these biases, respectively, background points were not selected randomly but instead selected proportional to the human footprint as a measure of accessibility and IMR as a measure of health system performance and access. Finally, this study provides no description of the roles of individual species as reservoirs, whether operating as maintenance, amplification, or bridging hosts, since neither infection competence nor infection susceptibility was measured in these species in the current study. Rather, individual species’ landscape suitability was used to 1) calculate species richness and evaluate this as a landscape feature of importance to the distribution of KFDV outbreak occurrence, particularly with respect to forest loss, and 2) identify individual species that may serve as useful sentinels in the development of wildlife surveillance mechanisms. Therefore, rather than defining how mammalian community ecology determines KFDV infection ecology, the current work delineates optimal landscape targets for developing the surveillance instruments necessary to appropriately monitor and sample wildlife so that the necessary community ecology and infection ecology data that are needed can be generated.

## Conclusions

This study provides novel metrics of mammalian species richness in the Western Ghats, one of the world’s critical biodiversity hotspots, and demonstrates the influence of this landscape feature and its interaction with deforestation on the expansion of one of India’s most important emerging zoonotic arboviruses. The insight gained can provide municipally-directed targets for landscape-based One Health surveillance of animals and humans in the face of anthropogenic landscape change, which can begin to provide a definitive understanding of KFDV infection ecology while simultaneously identifying the wildlife-human interfaces most vulnerable to pathogen spillover and which may yield the highest benefit from tailored intervention.

## Author contributions

Conceptualization: Michael Walsh, Rashmi Bhat, Venkatesh Nagarajan-Radha, Shailendra Sawleshwarkar, Chiranjay Mukhopadhyay; Data curation: Michael Walsh; Formal analysis: Michael Walsh; Investigation: Michael Walsh, Rashmi Bhat, Venkatesh Nagarajan-Radha, Prakash Narayanan, Navya Vyas, Shailendra Sawleshwarkar, Chiranjay Mukhopadhyay; Methodology: Michael Walsh, Rashmi Bhat, Venkatesh Nagarajan-Radha, Shailendra Sawleshwarkar; Supervision: Michael Walsh, Chiranjay Mukhopadhyay; Validation: Michael Walsh, Rashmi Bhat, Venkatesh Nagarajan-Radha, Shailendra Sawleshwarkar, Chiranjay Mukhopadhyay; Visualization: Michael Walsh; Writing - original draft: Michael Walsh, Rashmi Bhat, Venkatesh Nagarajan-Radha, Prakash Narayanan, Navya Vyas, Shailendra Sawleshwarkar, Chiranjay Mukhopadhyay; Writing - Michael Walsh, Rashmi Bhat, Venkatesh Nagarajan-Radha, Prakash Narayanan, Navya Vyas, Shailendra Sawleshwarkar, Chiranjay Mukhopadhyay.

