## Supplementary material for "Low mammalian species richness is associated with Kyasanur Forest Disease outbreak risk in deforested landscapes in the Western Ghats, India"

S1 Table 1. Species distribution ensemble models based on the weighted means of boosted regression tree-, random forest-, and generalised additive model-based species distributions. Performance is indicated by the area under the receiver operating characteristic curve (AUC) and fit is indicated by the deviance. The true skill statistic was used to mark the cutoff of the estimated species distribution function demarcating species presence.

| **Mammal species** | **Observations** | **AUC (%)** | **TSS** | **Deviance** |
|---|---|---|---|---|
| *Axis axis* | 185 | 93 | 0.80 | 0.57 |
| *Macaca radiata* | 359 | 88 | 0.69 | 0.88 |
| *Funambulus palmarum* | 239 | 90 | 0.69 | 0.77 |
| *Ratufa indica* | 217 | 95 | 0.88 | 0.49 |
| *Bos gaurus* | 153 | 92 | 0.76 | 0.69 |
| *Elephas maximus* | 145 | 96 | 0.85 | 0.47 |
| *Pteropus giganteus* | 138 | 94 | 0.81 | 0.61 |
| *Panthera tigris* | 106 | 95 | 0.81 | 0.52 |
| *Sus scrofa* | 108 | 88 | 0.69 | 0.75 |
| *Rusa unicolor* | 97 | 97 | 0.82 | 0.50 |
| *Gazella bennettii* | 88 | 100 | 0.98 | 0.36 |
| *Semnopithecus hypoleucos* | 92 | 86 | 0.66 | 0.93 |
| *Semnopithecus priam* | 56 | 87 | 0.65 | 0.87 |
| *Funambulus tristriatus* | 50 | 96 | 0.90 | 0.51 |
| *Semnopithecus johnii* | 50 | 91 | 0.87 | 0.65 |
| *Semnopithecus dussumieri* | 48 | 87 | 0.64 | 0.87 |
| *Funambulus pennantii* | 43 | 89 | 0.71 | 0.78 |
| *Antilope cervicapra* | 42 | 83 | 0.60 | 0.91 |
| *Canis aureus* | 52 | 85 | 0.66 | 0.92 |
| *Cuon alpinus* | 40 | 95 | 0.90 | 0.50 |
| *Macaca silenus* | 34 | 81 | 0.61 | 0.87 |
| *Panthera pardus* | 34 | 77 | 0.50 | 0.99 |
| *Melursus ursinus* | 33 | 90 | 0.78 | 0.67 |
| *Lepus nigricollis* | 32 | 84 | 0.67 | 0.88 |

S2 Figure 1. Individual species distributions.

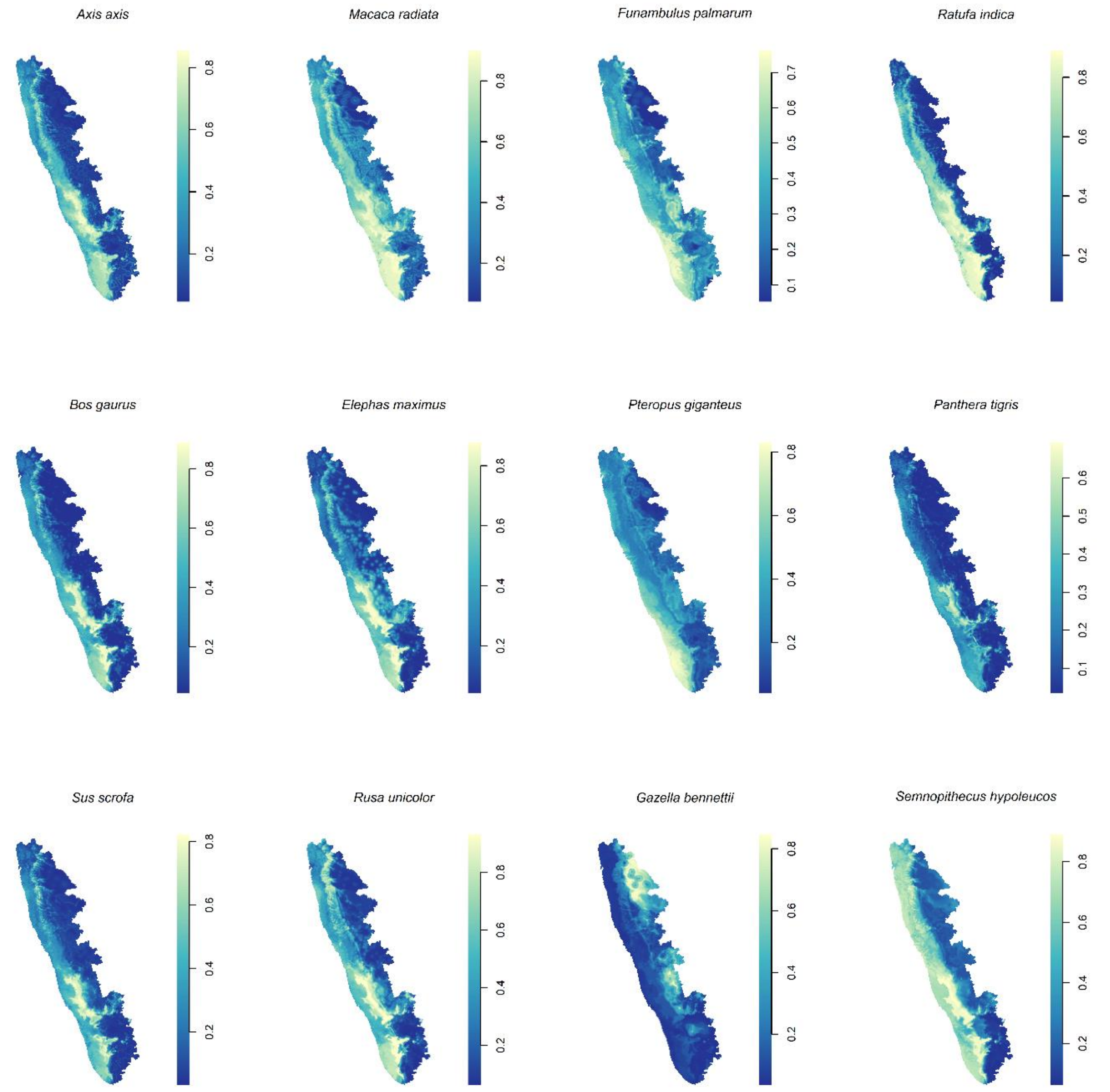

S3 Figure 2. Individual species distributions.

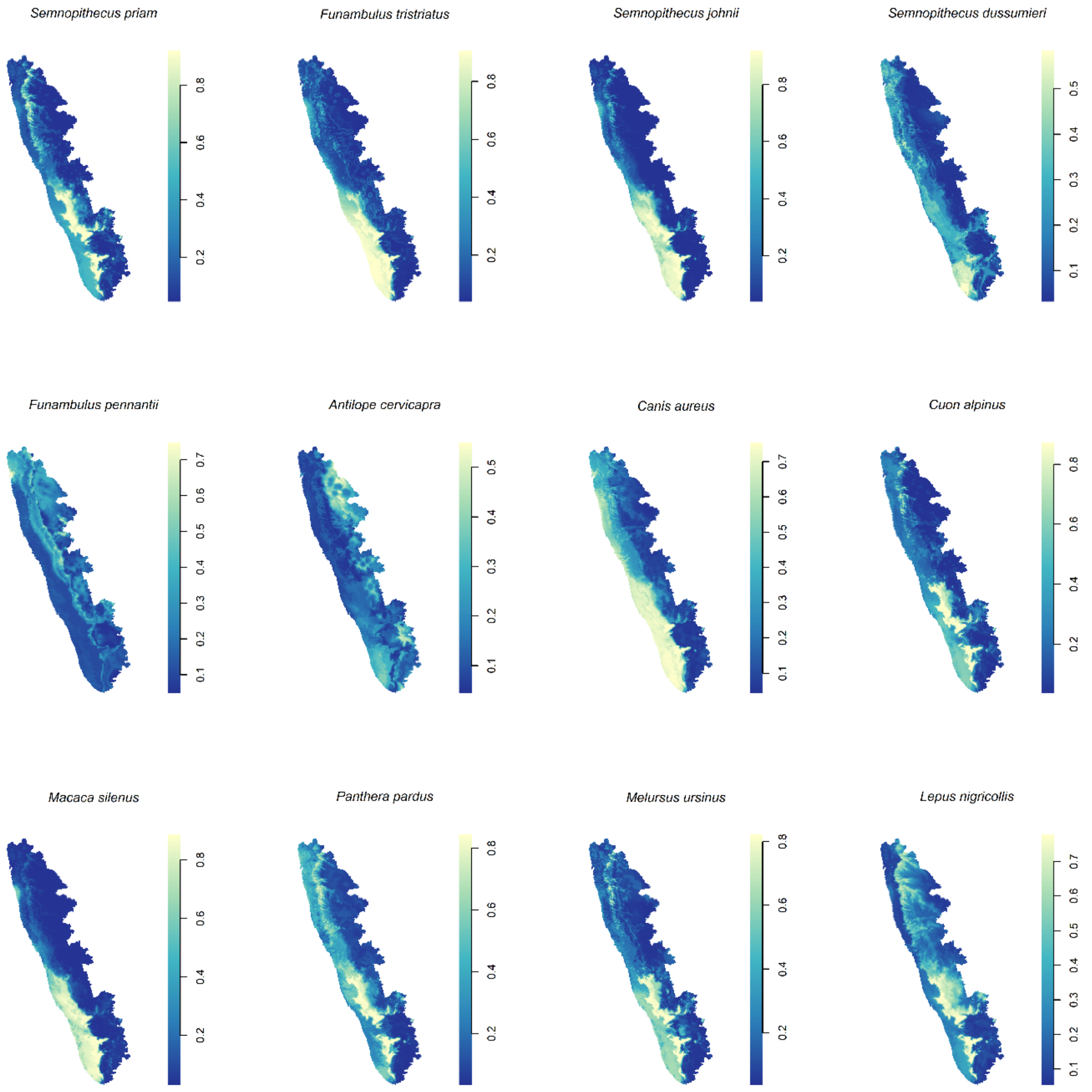

S4 Table 2. Secondary analysis species distribution ensemble models based on the weighted means of boosted regression tree-, random forest-, and generalised additive model-based species distributions. These models used an alternate India-wide sample of mammalian observations for a sensitivity analysis comparison of species richness estimates based on regional (S1 Table 1) and country-wide samples. Performance is indicated by the area under the receiver operating characteristic curve (AUC) and fit is indicated by the deviance. The true skill statistic was used to mark the cutoff of the estimated species distribution function demarcating species presence.

| **Mammal species** | **Observations** | **AUC (%)** | **TSS** | **Deviance** |
|---|---|---|---|---|
| *Axis axis* | 626 | 96 | 0.81 | 0.57 |
| *Macaca radiata* | 360 | 89 | 0.67 | 0.80 |
| *Funambulus palmarum* | 252 | 91 | 0.73 | 0.74 |
| *Ratufa indica* | 257 | 97 | 0.91 | 0.49 |
| *Bos gaurus* | 253 | 96 | 0.81 | 0.53 |
| *Elephas maximus* | 252 | 98 | 0.89 | 0.43 |
| *Pteropus giganteus* | 242 | 88 | 0.61 | 0.88 |
| *Panthera tigris* | 339 | 95 | 0.80 | 0.59 |
| *Sus scrofa* | 351 | 87 | 0.69 | 0.88 |
| *Rusa unicolor* | 297 | 93 | 0.74 | 0.71 |
| *Gazella bennettii* | 159 | 90 | 0.70 | 0.77 |
| *Semnopithecus hypoleucos* | 92 | 86 | 0.66 | 0.93 |
| *Semnopithecus priam* | 56 | 87 | 0.65 | 0.87 |
| *Funambulus tristriatus* | 50 | 96 | 0.90 | 0.51 |
| *Semnopithecus johnii* | 50 | 91 | 0.87 | 0.65 |
| *Semnopithecus dussumieri* | 330 | 91 | 0.70 | 0.74 |
| *Funambulus pennantii* | 431 | 82 | 0.58 | 0.99 |
| *Antilope cervicapra* | 171 | 89 | 0.80 | 0.74 |
| *Canis aureus* | 322 | 88 | 0.68 | 0.83 |
| *Cuon alpinus* | 61 | 91 | 0.70 | 0.77 |
| *Macaca silenus* | 34 | 81 | 0.61 | 0.87 |
| *Panthera pardus* | 108 | 82 | 0.57 | 1.03 |
| *Melursus ursinus* | 64 | 85 | 0.60 | 0.95 |
| *Lepus nigricollis* | 66 | 65 | 0.40 | 1.33 |
| *Boselaphus tragocamelus* | 398 | 93 | 0.80 | 0.64 |
| *Herpestes edwardsii* | 55 | 79 | 0.54 | 1.08 |
| *Paradoxurus hermaphroditus* | 48 | 64 | 0.44 | 1.24 |
| *Vulpes bengalensis* | 40 | 77 | 0.54 | 1.01 |
| *Hyaena hyaena* | 39 | 70 | 0.50 | 0.89 |
| *Lutrogale perspicillata* | 30 | 86 | 0.72 | 0.76 |

S5 Figure 3. Individual species distributions estimated with alternate country-wide sample.

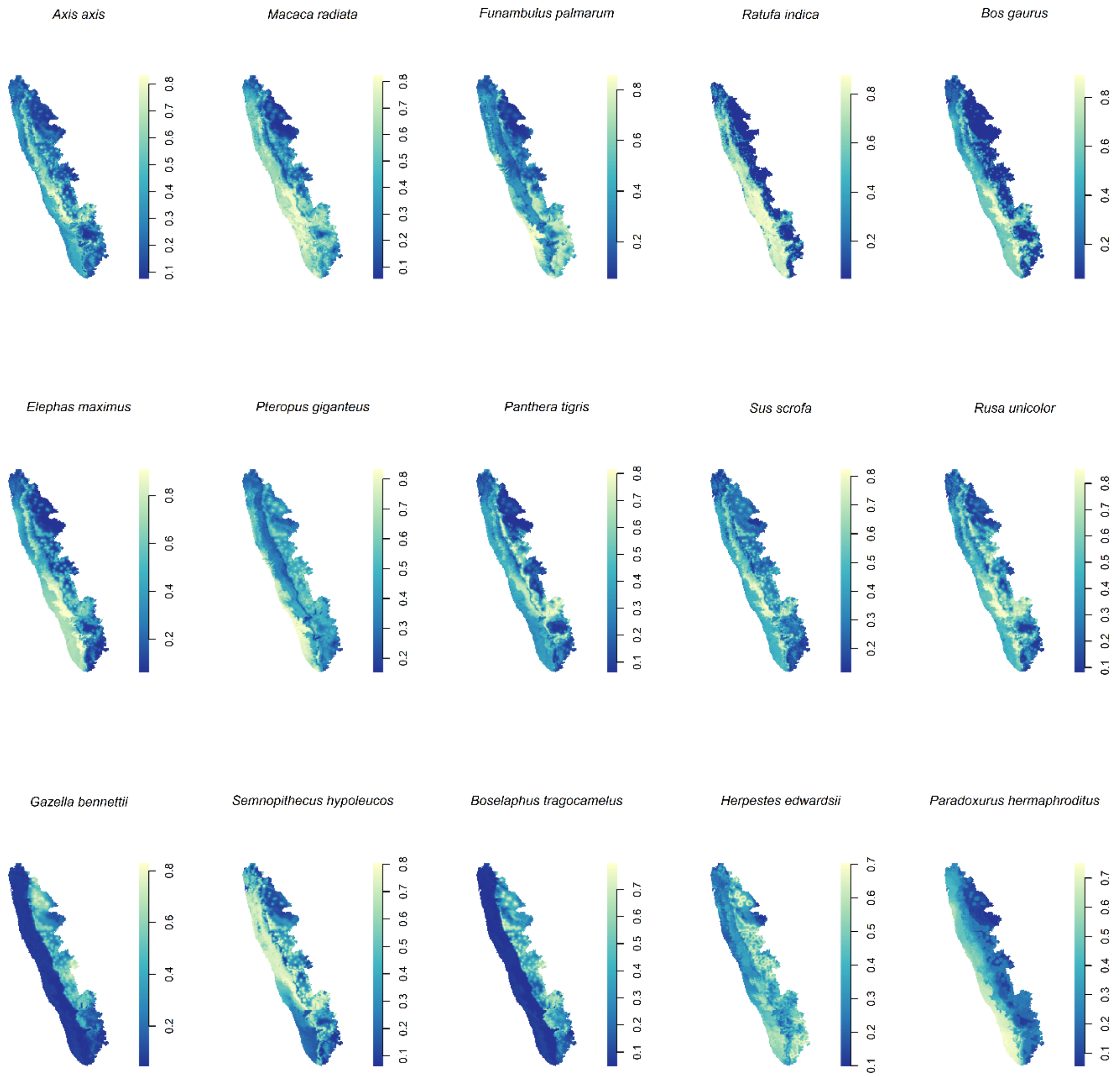

S6 Figure 4. Individual species distributions estimated with alternate country-wide sample.

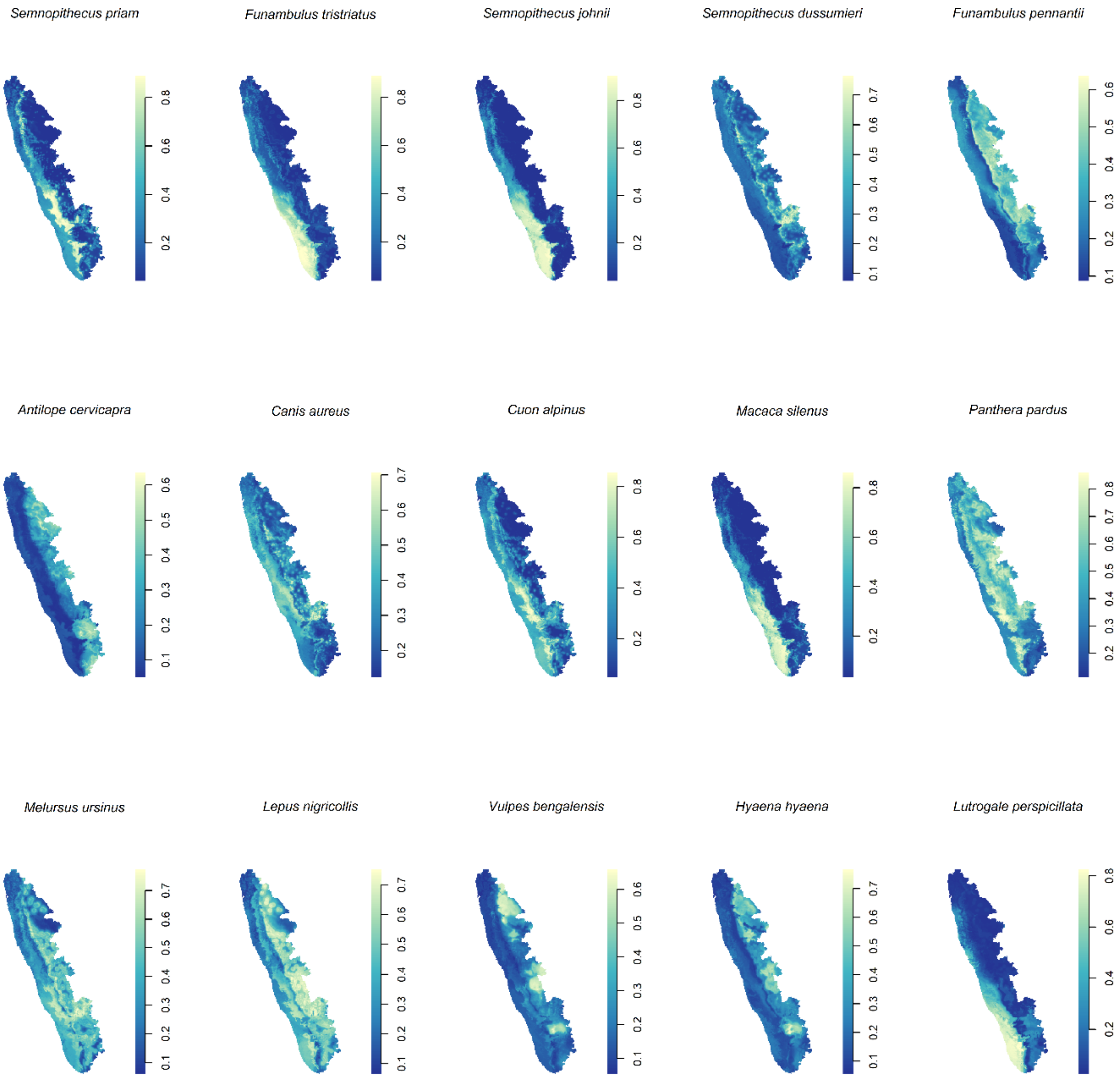

S7 Table 3. Regression coefficients and 95% confidence intervals for the associations between Kyasanur Forest disease virus outbreaks and landscape features as derived from inhomogeneous Poisson models of the outbreak point process and using the alternate country-wide sample to estimate species richness. Coefficients represent adjusted associations wherein each landscape feature is adjusted for all other features included in the multiple point process model. The Akaike information criterion (AIC) and area under the receiver operating characteristic curve (AUC) show model fit and performance, respectively.

| **Landscape features** | **Coefficient** | **95% confidence interval** | **AIC** | **AUC** |
|---|---|---|---|---|
| Mammal richness | 0.59 | 0.25 – 0.93 | -5.09 | 0.71 |
| Forest loss (deciles) | 4.99 | 2.51 – 7.45 | | |
| Mammal richness:forest loss interaction | -0.35 | -0.55 – -0.16 | | |
| Annual precipitation (100 mm) | 0.04 | 0.01 – 0.06 | | |
| Annual temperature (Celsius) | -0.01 | -0.02 – 0.001 | | |

S8 Figure 5. The distribution of Kyasanur Forest disease virus outbreak risk derived from the inhomogeneous Poisson model, and using the alternate country-wide sample to estimate species richness, is presented in the centre panel with the lower and upper 95% confidence limits to the left and right, respectively.

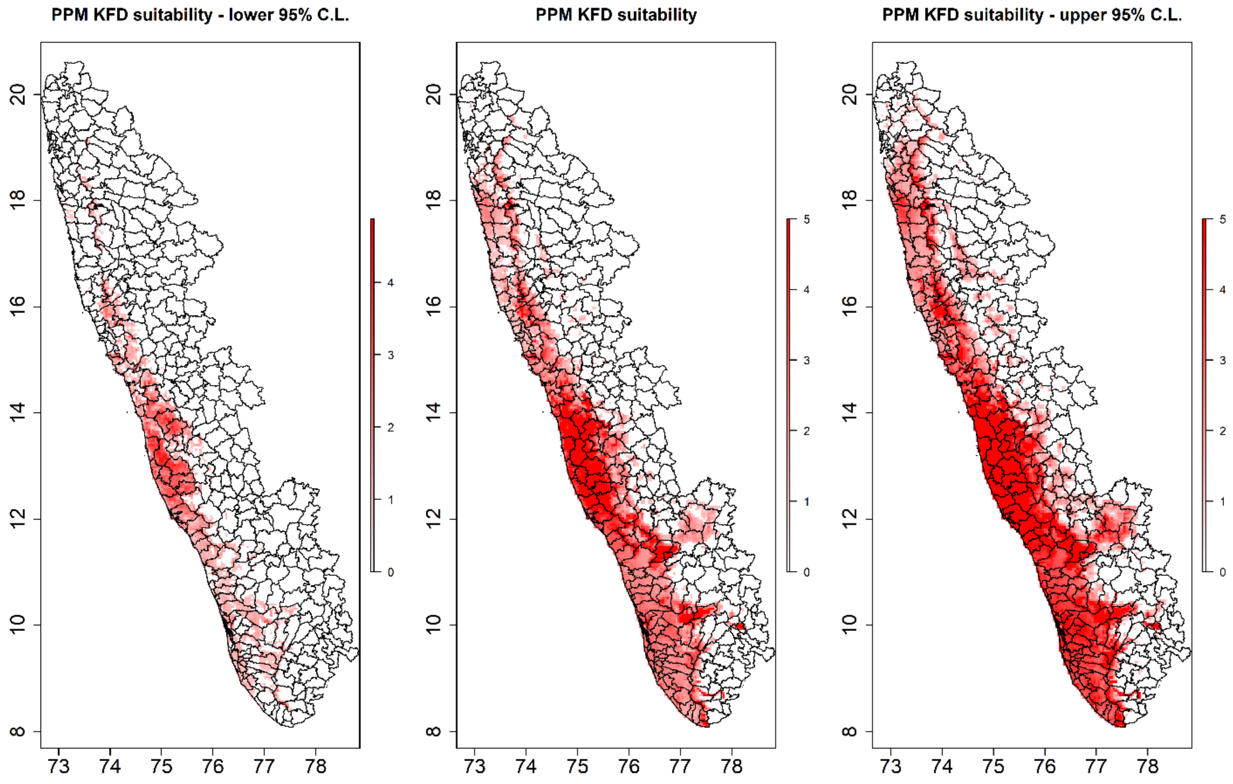

S9 Table 4. Taluk-level multiple integrated nested Laplace approximation model (zero-inflated Poisson family) of Kyasanur Forest disease virus outbreaks. Mammal species richness was adjusted for the biotic constraints of sympatric species using the SESAM framework. The Watanabe-Akaike information criterion (WAIC) was used to assess model fit.

| **Landscape features** | **Coefficient** | **95% credible interval** | **WAIC** |
|---|---|---|---|
| Mammal richness$_{SESAM}$ | 1.51 | 0.40 – 2.63 | 215.16 |
| Forest loss (deciles) | 8.32 | 2.55 – 13.88 | |
| Mammal richness:forest loss interaction | -0.80 | -1.39 – -0.20 | |
| Annual precipitation (100 mm) | 0.10 | 0.01 – 0.20 | |
| Annual temperature (Celsius) | 0.003 | -0.045 – 0.044 | |

S10 Figure 6. The distribution of Kyasanur Forest disease virus suitability derived from the integrated nested Laplace approximation model (zero-inflated Poisson family) is presented in the centre panel with the lower and upper 95% credible limits to the left and right, respectively.

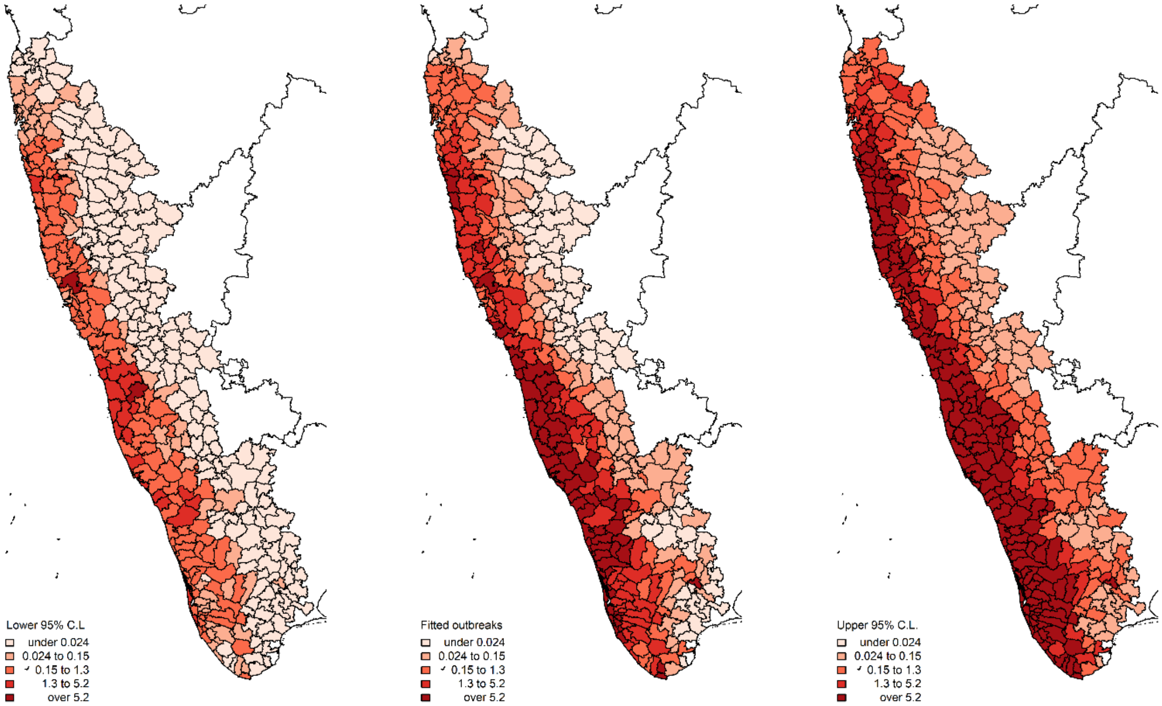

S11 Table 5. Taluk-level integrated nested Laplace approximation model (binomial family) of Kyasanur Forest disease virus outbreaks using the alternate country-wide sample to estimate species richness. Mammal species richness was adjusted for the biotic constraints of sympatric species using the SESAM framework. The Watanabe-Akaike information criterion (WAIC) was used to assess model fit.

| **Landscape features** | **Coefficient** | **95% credible interval** | **WAIC** |
|---|---|---|---|
| Mammal richness$_{SESAM}$ | 1.25 | 0.48 – 2.08 | 151 |
| Forest loss (deciles) | 8.00 | 1.37 – 14.50 | |
| Mammal richness:forest loss interaction | -0.75 | -1.34 – -0.18 | |
| Annual precipitation (100 mm) | 0.00 | 0.00 – 0.1 | |
| Annual temperature (Celsius) | -0.02 | -0.038 – -0.001 | |

S12 Figure 7. The adjusted distribution of mammalian species richness among the Western Ghats using the alternate country-wide sample. Species richness was adjusted using the SESAM framework to introduce biotic constraints to adjust the presence of each species for the presence of other species within each taluk.

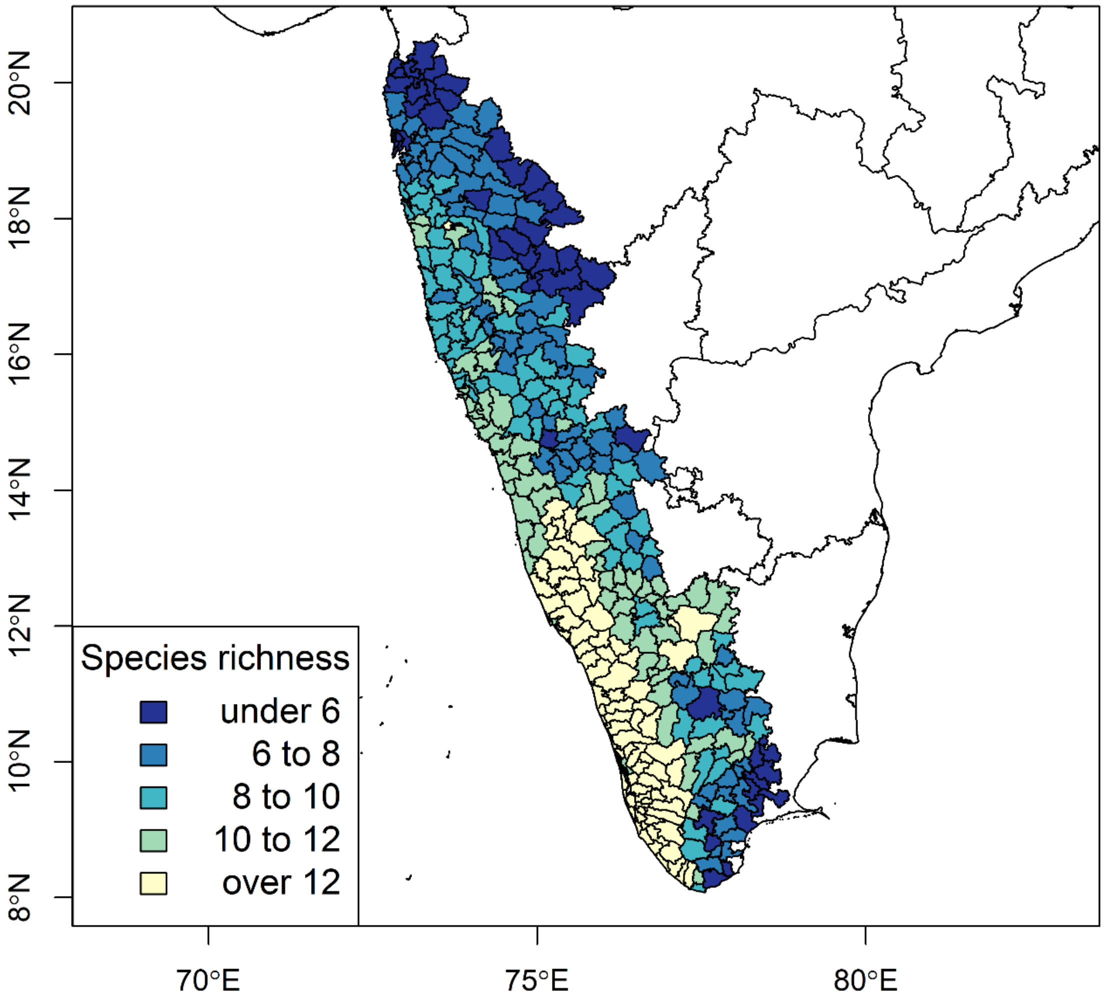

S13 Table 6. Unadjusted regression coefficients and 95% confidence intervals for the crude associations between Kyasanur Forest disease virus outbreaks and individual mammal species as derived from inhomogeneous Poisson models of the outbreak point process. The Akaike information criterion (AIC) and area under the receiver operating characteristic curve (AUC) show model fit and performance, respectively.

| Mammal species | Coefficient | 95% confidence interval | AIC | AUC |
|---|---|---|---|---|
| *Axis axis* | 4.17 | 2.87 – 5.49 | 20.54 | 0.70 |
| *Macaca radiata* | 3.60 | 2.25 – 4.96 | 32.82 | 0.65 |
| *Funambulus palmarum* | 3.63 | 2.03 – 5.22 | 43.92 | 0.61 |
| *Ratufa indica* | 2.64 | 1.52 – 3.76 | 13.07 | 0.62 |
| *Bos gaurus* | 3.02 | 1.95 – 4.09 | 33.89 | 0.67 |
| *Elephas maximus* | 2.59 | 1.51 – 3.66 | 42.69 | 0.61 |
| *Pteropus giganteus* | 0.44 | -1.01 – 1.89 | 64.14 | 0.53 |
| *Panthera tigris* | 1.76 | -0.06 – 3.59 | 61.15 | 0.63 |
| *Sus scrofa* | 3.31 | 2.14 – 4.48 | 36.06 | 0.67 |
| *Rusa unicolor* | 2.84 | 1.74 – 3.94 | 37.57 | 0.64 |
| *Gazella bennettii* | -4.97 | -8.03 – -1.90 | 46.85 | 0.58 |
| *Semnopithecus hypoleucos* | 5.54 | 3.68 – 7.41 | 4.73 | 0.70 |
| *Semnopithecus priam* | 2.57 | 1.66 – 3.48 | 37.19 | 0.65 |
| *Funambulus tristriatus* | 0.18 | -0.76 – 1.12 | 64.34 | 0.59 |
| *Semnopithecus johnii* | 0.76 | -0.10 – 1.62 | 61.67 | 0.67 |
| *Semnopithecus dussumieri* | 2.52 | 0.72 – 4.32 | 57.46 | 0.74 |
| *Funambulus pennantii* | -9.71 | -14.74 – -4.68 | 42.00 | 0.62 |
| *Antilope cervicapra* | -11.68 | -17.13 – -6.22 | 35.94 | 0.50 |
| *Canis aureus* | 2.41 | 1.29 – 3.53 | 46.12 | 0.67 |
| *Cuon alpinus* | 1.63 | 0.58 – 2.69 | 56.28 | 0.65 |
| *Macaca silenus* | 0.45 | -0.50 – 1.39 | 63.65 | 0.67 |
| *Panthera pardus* | 3.52 | 2.38 – 4.66 | 27.90 | 0.67 |
| *Melursus ursinus* | 1.14 | 0.003 – 2.28 | 60.89 | 0.65 |
| *Lepus nigricollis* | 2.73 | 1.49 – 3.98 | 47.72 | 0.56 |

S14 Table 7. Taluk-level regression coefficients and 95% credible intervals for the associations between Kyasanur Forest disease virus outbreaks and each species abundance (per 10 species-present pixels per taluk) and relative abundance as derived from integrated nested Laplace approximation models (binomial family) of Kyasanur Forest Disease virus outbreaks. Coefficients represent crude associations from bivariate models.

| **Mammal species abundance/relative abundance** | **Coefficient** | **95% credible interval** | **WAIC** |
|---|---|---|---|
| *Axis axis* abundance | 0.02 | 0.00 – 0.04 | 162.74 |
| *Axis axis* relative abundance | ***29.09*** | ***3.84 – 51.45*** | ***159.07*** |
| *Macaca radiata* abundance | ***0.01*** | ***0.015 – 0.02*** | ***155.13*** |
| *Macaca radiata* relative abundance | -0.035 | -2.85 – 2.20 | 164.55 |
| *Funambulus palmarum* abundance | 0.01 | -0.02 – 0.03 | 165.39 |
| *Funambulus palmarum* relative abundance | -1.17 | -6.37 – 2.61 | 164.39 |
| *Ratufa indica* abundance | -0.01 | -0.10 – 0.05 | 164.82 |
| *Ratufa indica* relative abundance | -10.08 | -60.53 – 32.34 | 163.04 |
| *Bos gaurus* abundance | 0.01 | -0.01 – 0.02 | 164.05 |
| *Bos gaurus* relative abundance | 3.84 | -10.99 – 15.75 | 164.12 |
| *Elephas maximus* abundance | 0.03 | -0.03 – 0.07 | 163.88 |
| *Elephas maximus* relative abundance | 12.16 | -37.16 – 55.51 | 163.19 |
| *Pteropus giganteus* abundance | -0.02 | -0.09 – 0.02 | 163.96 |
| *Pteropus giganteus* relative abundance | -17.76 | -47.49 – 3.83 | 160.01 |
| *Panthera tigris* abundance | 0.00 | -62.09 – 62.03 | 162.69 |
| *Panthera tigris* relative abundance | 0.00 | -62.09 – 62.03 | 162.69 |
| *Sus scrofa* abundance | 0.01 | -0.01 – 0.02 | 163.80 |
| *Sus scrofa* relative abundance | 12.22 | -7.62 – 28.79 | 162.97 |
| *Rusa unicolor* abundance | 0.01 | 0.00 – 0.02 | 162.88 |
| *Rusa unicolor* relative abundance | -0.20 | -6.44 – 4.39 | 164.90 |
| *Gazella bennettii* abundance | -0.01 | -0.03 – 0.01 | 178.54 |
| *Gazella bennettii* relative abundance | -10.34 | -53.79 – 18.15 | 175.72 |
| *Semnopithecus hypoleucos* abundance | ***0.02*** | ***0.01 – 0.03*** | ***139.72*** |
| *Semnopithecus hypoleucos* relative abundance | ***2.24*** | ***0.98 – 3.53*** | ***152.75*** |
| *Semnopithecus priam* abundance | ***0.01*** | ***0.001 – 0.02*** | ***160.23*** |
| *Semnopithecus priam* relative abundance | 5.64 | -0.40 – 11.31 | 161.87 |
| *Funambulus tristriatus* abundance | -0.01 | -0.06 – 0.02 | 164.13 |
| *Funambulus tristriatus* relative abundance | -17.00 | -49.30 – 7.78 | 161.69 |
| *Semnopithecus johnii* abundance | 0.00 | -0.01 – 0.01 | 164.51 |
| *Semnopithecus johnii* relative abundance | -6.22 | -22.30 – 6.67 | 163.55 |
| *Semnopithecus dussumieri* abundance | -3.01 | -40.37 – 19.42 | 163.68 |
| *Semnopithecus dussumieri* relative abundance | -3.01 | -40.37 – 19.42 | 163.68 |
| *Funambulus pennantii* abundance | -0.82 | -28.21 – 14.46 | 170.30 |
| *Funambulus pennantii* relative abundance | -9.05 | -55.73 – 22.27 | 163.78 |
| *Antilope cervicapra* abundance | -0.02 | -0.08 – 0.03 | 179.59 |
| *Antilope cervicapra* relative abundance | -13.24 | -63.62 – 22.42 | 166.38 |
| *Canis aureus* abundance | 0.00 | -0.02 – 0.01 | 166.73 |
| *Canis aureus* relative abundance | -4.86 | -12.21 – 1.00 | 162.24 |
| *Cuon alpinus* abundance | 0.00 | -0.02 – 0.02 | 164.38 |
| *Cuon alpinus* relative abundance | 6.17 | -20.42 – 27.85 | 164.18 |
| *Macaca Silenus* abundance | 0.00 | -0.01 – 0.01 | 165.03 |
| *Macaca Silenus* relative abundance | -5.17 | -15.02 – 2.76 | 163.12 |
| *Panthera pardus* abundance | ***0.02*** | ***0.01 – 0.03*** | ***147.76*** |
| *Panthera pardus* relative abundance | ***5.06*** | ***1.79 – 8.25*** | ***156.37*** |
| *Melursus ursinus* abundance | 0.01 | -0.01 – 0.02 | 163.44 |
| *Melursus ursinus* relative abundance | 11.54 | -4.69 – 25.33 | 162.64 |
| *Lepus nigricollis* abundance | 0.01 | 0.00 – 0.02 | 162.10 |
| *Lepus nigricollis* relative abundance | 2.26 | -8.24 – 10.70 | 164.28 |

S15 Figure 8. Taluk-level species abundance and relative abundance for primate species associated with KFDV outbreaks (S14 Table 7).

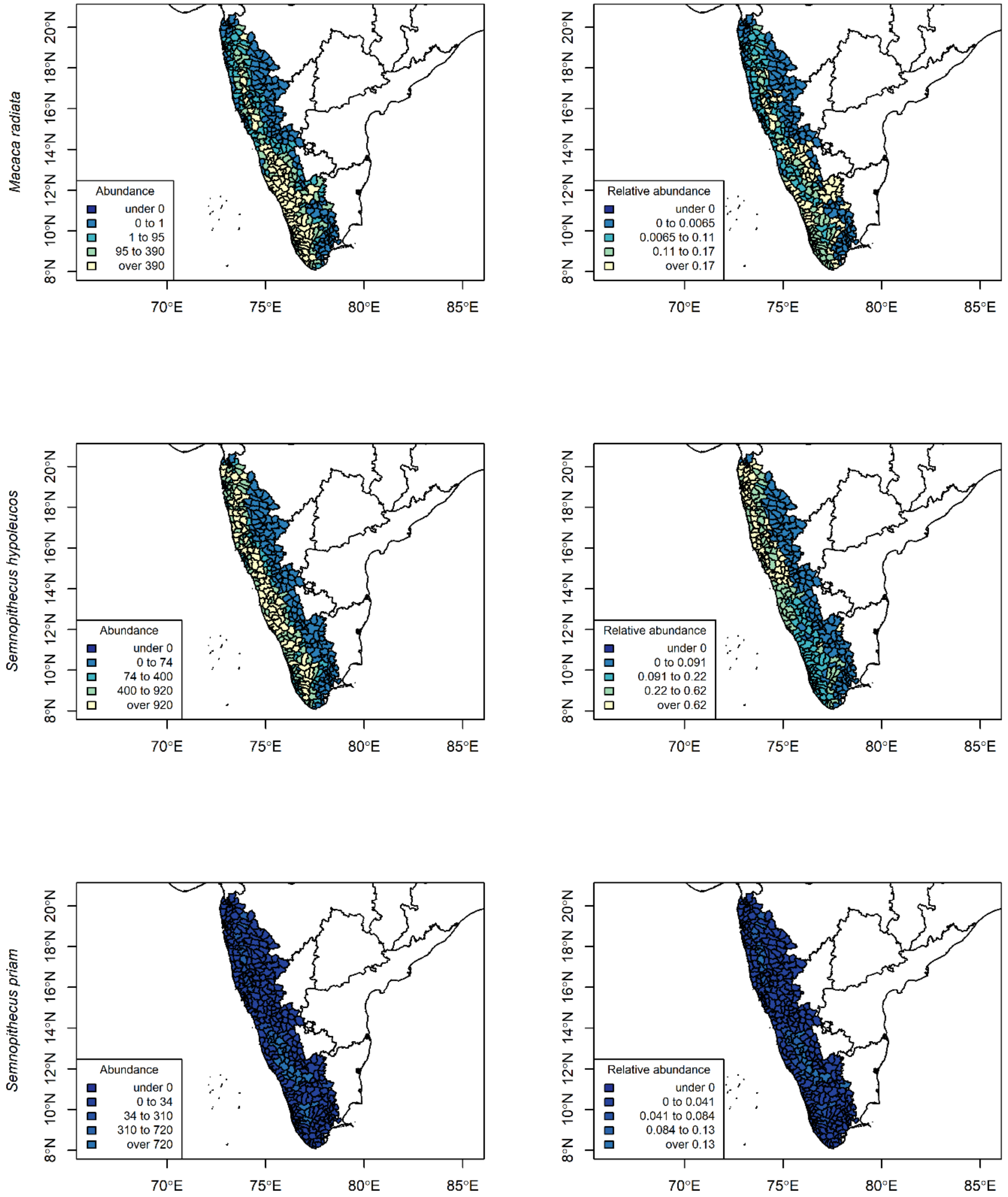
